# Evaluation of rice wild relatives as a source of traits for adaptation to iron toxicity and enhanced grain quality

**DOI:** 10.1101/771352

**Authors:** Birgit Bierschenk, Melle Tilahun Tagele, Basharat Ali, Md Ashrafuzzaman, Lin-Bo Wu, Matthias Becker, Michael Frei

**Author notes:** These authors contributed equally to the work (Birgit Bierschenk, Melle Tilahun Tagele). Corresponding author: INRES, University of Bonn, Katzenburgweg 5, 53115 Bonn.

## Abstract

Rice wild relatives (RWR) constitute an extended gene pool that can be tapped for the breeding of novel rice varieties adapted to abiotic stresses such as iron (Fe) toxicity. Therefore, we screened 75 *Oryza* genotypes including 16 domesticated *O. sativas*, one *O. glaberrima*, and 58 RWR representing 21 species, for tolerance to Fe toxicity. Plants were grown in a semi-artificial greenhouse setup, in which they were exposed either to control conditions, an Fe shock during the vegetative growth stage (acute treatment), or to a continuous moderately high Fe level (chronic treatment). In both stress treatments, foliar Fe concentrations were characteristic of Fe toxicity, and plants developed foliar stress symptoms, which were more pronounced in the chronic Fe stress especially toward the end of the growing season. Among the genotypes that produced seeds, only the chronic stress treatment significantly reduced yields due to increases in spikelet sterility. Moreover, a moderate but non-significant increase in grain Fe concentrations, and a significant increase in grain Zn concentrations were seen in chronic stress. Both domesticated rice and RWR exhibited substantial genotypic variation in their responses to Fe toxicity. Although no RWR strikingly outperformed domesticated rice in Fe toxic conditions, some genotypes scored highly in individual traits. Two *O. meridionalis* accessions were best in avoiding foliar symptom formation in acute Fe stress, while an *O. rufipogon* accession produced the highest grain yields in both chronic and acute Fe stress. In conclusion, this study provides the basis for using interspecific crosses for adapting rice to Fe toxicity.

## Introduction

Crop wild relatives (CWR) are widely regarded as a rich source of genetic variation for the improvement of domesticated crops, which can be tapped more effectively than ever before due to the progress in CWR genomics [1,2]. Rice is one of the most important cereal crops in the world in terms of annual production and provides the staple food for around half of the world’s population [3]. Two domesticated rice species are widely grown: *Oryza sativa* L., which was domesticated from the wild progenitors *O. rufipogon* and *O. nivara* in Asia around 10, 000 years ago [4,5] and *Oryza glaberrima* Steud., which was independently domesticated from the wild progenitor *Oryza barthii* in West Africa around 3000 years ago [6]. In addition, archeological evidence suggest that rice wild relatives (RWR) originating from South America, such as *O. glumaepatula* underwent a process of domestication in the mid-Holocene by Native South Americans, but were abandoned later on when immigrants from Europe changed the fate of the continent and its inhabitants to a great extent [7]. The whole *Oryza* genus, which can be regarded as a primary, secondary and tertiary gene pool for rice breeding, contains 24 species [8]. It can be further classified into four complexes of closely related species with the same genome groups, between which fertile crossing is possible [9,10]. These are the *O. sativa* complex comprising the diploid AA species (Supplementary Table S1), the *O. officinalis* complex, the *O. granulata* complex and the *O. ridleyi* complex [11]. In addition *O. sativa f. spontanea* or ‘weedy rice’ can be found in rice growing regions all over the world [12] and likely represents a hybrid between cultivated rice varieties and the closely related RWR *O. rufipogon* or *O. nivara,* but it can also result from spontaneous mutations of *O. sativa* varieties or crosses between them [13].

Often, RWR are seen as weeds and thus as a major problem for rice cultivation, but they also bear a lot of potential, as *Oryza* species are distributed around the globe in very different environments, and their genome therefore might harbor adaptive genes to several biotic and abiotic stresses. Some of these adaptive traits were previously used to develop commercial varieties. Breeding of male sterile rice lines, which is required for the commercial production of hybrids, was originally only possible with the cytoplasm of *O. sativa* f. *spontanea* [14], and those hybrids were still planted on nearly half of the rice planting area of China in 2006 [15]. In 1974 the first rice varieties containing resistance genes to the grassy stunt virus from *Oryza nivara* were released [14]. Crosses with *O. officinalis* lead to cultivars with resistance to brown plant hoppers, of which three were released in Vietnam [14]. Several other resistances of RWR against pests, pathogens and abiotic stresses and even yield improving traits have been identified but not yet been taken advantage of [8,9,14]. In some cases fertile crosses with *O. sativa* were accomplished. A resistance gene against bacterial blight from *O. longistaminata* was crossed into popular varieties [16], although the progenies were not established as commercial varieties. Crosses with the remote relative *O. brachyantha* were successful, even though the desired resistance to yellow stem borer could not be preserved [17]. Abiotic stresses constitute another category of yield constraints that may be mitigated by the use of RWR in breeding. *O. officinalis* has been proven as useful to address heat stress, because in a cross with *O. sativa* early-morning flowering lead to higher fertility due to the avoidance of high midday temperatures [18]. A cold tolerant *O. rufipogon* genotype contained several interesting resistance loci [19]. *O. coarctata* is used as a model to understand salt tolerance in rice, and successful crosses with the modern rice variety IR56 exist [8,20]. Aluminum tolerance in acidic soils from *O. rufipogon* was previously dissected into quantitative trait loci [21]. Other potential fields of application of RWR in tolerance breeding include drought and submergence [8]. Up to date not much is known about the chemical and nutritional grain composition of RWR and genetic factors that might be of use for grain quality breeding, although this approach may hold great potential [22].

Fe toxicity is one of the major mineral disorders affecting rice production, especially in parts of West Africa, Southeast Asia, Madagascar and the South of Brazil [23]. It frequently occurs because rice-growing soils are inherently rich in Fe, but more importantly the flooding of paddy soils leads to microbial reduction processes that release abundant soluble Fe^2+^ (ferrous Fe) [23,24]. When excessive Fe is taken up into rice plants it leads to the formation of oxidative stress via the Fenton reaction [25], visible leaf bronzing symptoms [26], and eventually yield losses due to reduced growth and increased spikelet sterility [27,28]. Fe toxicity either occurs as an acute stress during the vegetative stage, when abundant amounts of Fe are mobilized from adjacent slopes due to heavy rainfall, or as a chronic stress with a more gradual build-up of high Fe concentrations in soil solution [23,28]. As farmers have very few management options to address Fe toxicity, the breeding of Fe tolerant rice varieties represents the most promising strategy of adapting rice production to high Fe soils. A large number of genetic screening and genome mapping experiments have been conducted in the past to identify donors and loci that can be used for the breeding of tolerant germplasm [26,29,30]. These studies have revealed that Fe tolerance is a complex trait controlled by a large number of small and medium effect loci rather than large effect quantitative trait loci [26,30]. Moreover, the progress in breeding is hampered by limited consistency between different screening experiments and lack of genetic variation for certain desirable traits, such as the ability to translocate large amounts of Fe to the grains despite Fe toxicity [28]. Another limitation is that to date no RWR have been evaluated regarding their potential to contribute to Fe tolerance breeding.

Therefore, the present study was undertaken to explore traits of RWR that could be exploited for the breeding of Fe tolerant rice. Due to the recent report of genome sequences for thirteen RWR [5], and the availability of mapping populations derived from interspecific crosses of rice (e.g. [31] this is a timely approach that could lead to the fast discovery of novel genes for rice breeding and thus contribute to diversification and adaptation. Specifically, we hypothesized that (i) the variation observed in RWR for adaptation to chronic and acute Fe stress may exceed the variation observed in domesticated rice; (ii) despite low agronomic value RWR may possess specific traits that can be used in adaptive breeding; and (iii) when grown in high Fe conditions RWR may possess grain quality traits that differ from those observed in domesticated rice.

## Materials and Methods

### Plant material

Seeds of 60 rice accessions were provided by the International Rice Research Institute (IRRI) in Los Baños, the Philippines. They consisted of 20 wild species of the *Oryza* genus with different countries of origin, one cross of *O. glaberrima* and *O. barthii* and two *O. sativa* varieties, for which crosses with wild relatives have been made (IRRI 154 and Curinga). In addition, 15 accessions of cultivated rice species (*O. sativa* and one *O. glaberrima*) that had previously been screened in Fe toxicity experiments [26,32,33] were added from the seed stocks of the Institute of Crop Science and Resource Conservation (INRES).

### Germination

Where necessary, seeds dormancy was broken by placing seeds in an oven at 50°C for three days. Thirty seeds of each accession were then placed on a floating mesh in a germination box with about two liters of deionized water, making sure that the grains were not fully covered by the water. The seeds were incubated in darkness at a temperature of 33°C. After three days the first seeds germinated and were transferred to a germination tray floating on deionized water containing 10 μM Ferric sodium EDTA (Fe-Na-EDTA) and 0.5 mM calcium chloride (CaCl). The box was placed in a greenhouse until the seedlings were about 8 cm high. Solutions were exchanged every two days.

### Experimental setup and growing conditions

The experiment was conducted in a greenhouse at Campus Klein-Altendorf, which is an agricultural trial station of the University of Bon from May to October 2017. Six polders of 2 × 6 m dimension with a soil depth of about 50 cm were utilized for an experimental approach simulating acute and chronic iron toxicity as described previously [28]. Two independent polders each were used for the three treatments chronic iron stress, acute iron stress and control. Within each polder four sub-replicates containing all accessions were planted. The polders were ploughed and leveled manually and then kept flooded. As complete randomization was not recommendable due to unmanageable complexity, every accession was assigned a random number from 1 to 75 and this sequence was planted in a row. In every sub-replication the sequence started with a different number leading to a semi-randomized setup. Seedlings were planted with a spacing of 20 cm to each other and 10 cm to the borders. Every planting position was marked with a wooden stick to avoid confusions because of missing plants. Therefore, one polder consisted of 300 plants. In summary, the experiment was conducted as a two-factorial design (iron-treatment and genotype) with two true replicates per treatment (independent polders) and four semi-randomized sub-replicates in every polder.

The soil type in the polders was a clay silt with about 5 % smectite, 16 % vermiculite, 69 % illite and 10 % kaolinite, originating from a Luvisol on deep loess with an organic carbon content of 1.2 % and a pH between 6 and 7 [43]. Average day and night time temperatures were 28°C and 18.4°C respectively, the average relative humidity was 76.4% and the average CO_2_ concentration was 553 ppm.

All polders were watered regularly at least once a week. One hundred fifty g of potassium phosphate (K_2_HPO_4_) dissolved in water were applied to every polder. This corresponds to a concentration of about 6 g m^−2^ (60 kg ha^−1^) of potassium and 2.2 g m^−2^ (22 kg ha^−1^) of phosphorus. The same method was used to apply nitrogen in the form of urea. The application was split: 78 g were given five weeks after transplanting (WAT) together with the potassium phosphate, while another 78 g were applied nine WAT leading to a total application of 6 g m^−2^ (60 kg ha^−1^) of nitrogen. As some accessions began to lodge in the reproductive phase the plants were supported with tonkin sticks. Perforated polypropylene bags, so called crispac-bags (305 × 450 mm, perforation diameter 0.5 mm, vendor Baumann Saatzuchtbedarf GmbH, Waldenburg, Germany) were wrapped around ripening panicles to avoid seed shattering of wild rice accessions. The insecticide ‘Perfekthion’ (BASF, Ludwigshafen, Germany; active compound dimethoate) was sprayed on the plants due to an infestation with *Lissorhoptrus* ssp. 5 weeks after transplantation. Predatory mites (*Phytoseiulus persimilis* and *Amblyseiuscalifornicus*) were applied during the reproductive phase to control mite infestation.

### Iron treatment

Iron treatment was started six weeks after transplanting (WAT). A total of 10 kg of iron(II) sulfate heptahydrate (FeSO_4_×7H_2_O) was applied to the iron treatment polders. Five kg were applied six and seven WAT to obtain an approximate concentration of 1500 mg l^−1^ of reduced iron in the upper soil solution in acute polders, while 1 kg was given to chronic polders every week until 16 WAT to reach a concentration of 200 to 300 mg l^−1^ [28]. An even distribution of all the chemicals was ensured by diluting the powders in water and applying it with a watering can.

### Collection of data during growth

Leaf bronzing score was determined at one-week interval by visual observation of bronzing symptoms on the two youngest fully developed leaves of two tillers per plant. The percentage of leaf area affected by iron-induced symptoms was scored on a scale from 0-10 (in which 0 means no visual symptom and 10 indicates dead leaves) [26].

Leaves for iron analyses were sampled at 116 days after transplanting from genotypes selected based on contrasting symptom development at that stage. From each selected plant, the third youngest leaf was taken, dried, and subjected to Fe analyses as described below.

### Harvest

Harvest took place 24 and 27 WAT. In the first round all plants with mature panicles were harvested. Three weeks later, the remaining plants were harvested. Plants were cut close to the soil surface, the height was measured, the number of tillers and panicles was counted, the panicles were put into a crispac-bag and the straw biomass was folded to a bundle.

Straw biomass was dried at 60°C for at least 4 days and then weighed. The panicles were dried at 40°C for at least 4 days, then the seeds were detached from the spikes. Grain samples were weighed, and then two batches of 20 unfilled grains were counted and weighed. Unfilled grains were separated from the filled ones by winnowing and blowing. The filled grains were weighed and again two batches of 20 filled grains were counted and weighed. With this data the number of seeds per panicle, the thousand kernel weight (TKW) and the percentage of unfilled grains could be determined. In order to avoid errors due to the presence of awns in some accessions, two times 20 awns of three samples of all accessions with strongly developed awns were separated from grains, counted and weighed. The mean weight of awns was used to correct the weight calculations of these accessions as awns were removed along with the unfilled grains. The harvest index was calculated from the weight of filled grains in relation to the whole plant biomass in percent.

### Seed analysis

All RWR that produced seeds were included in the grain analyses, as well as the only *O. glaberrima*, and *Oryza sativa* Dom Sofid, IR 72, IRRI 154 and Curinga. Two samples per polder equaling four samples per treatment were chosen of every genotype. For dehusking of the seeds a Mixer Mill (MM 2000, manufacturer RETSCH, Haan, Germany) was used. The grinding jars made of polytetrafluorethen (PTFE) were filled to three quarters with sample material and different numbers of small (7 mm), medium (10 mm) and large (12 mm) grinding balls made of agate were used to ensure that the samples had no contact with Fe or Zn containing material. It was not possible to conduct the dehusking under standardized conditions as the seed properties differed widely between species and accession. Therefore, conditions were calibrated for every accession to protect the pericarp, aleurone layer and embryo as much as possible to produce brown rice. For grinding the agate balls were exchanged with one 20 mm PTFE Ball with a steel core, and all seeds were ground at speed 70 for one minute. All samples were dried again afterwards for 2 days at 60°C to ensure complete dryness.

Phytate analysis followed the procedure described by [34] and was adjusted to microplate format. For the Wade reagent, 18 mg of iron(III) chloride (FeCl_3_) and 350 mg of sulfosalicylic acid dihydrate (SSA · 2 H_2_O) were dissolved in 100 ml of deionized water. Of every sample of finely ground seeds, two times 50 mg were weighed into 2 mm Eppendorf-tubes to obtain two analytical replicates. After adding 1 ml of 3.5 % hydrochloric acid (HCl) all samples were stirred with a rotator (RS-RR 5, manufacturer Phoenix Instrument, Garbsen, Germany) at 40 rpm for 1 h at room temperature. Centrifugation of samples was carried out with a MIKRO 200 R (manufacturer Hettich, Buckinghamshire, England) at 21382 g for 10 minutes at 20°C. Two times 5 μl of every sample was each mixed with 145 μl of distilled water and 50 μl of wade reagent in a well-plate and the absorbance taken at 500 nm. The phytate concentration was calculated with a standard curve ranging from 0 to 10 μg phytate using the microplate analysis software Gen5 (developer BioTek, Winooski, VT, USA).

For iron and zinc determination in seeds, the ground material was first solubilized with a PDS-6 Pressure Digestion System (manufactured by Loftfields Analytical Solutions, NeuEichenberg, Germany). 250 mg ± 9 mg of every sample were weighed into one PTFE beaker each and the precise weight was noted. One blank and one standard of 250 mg MSC certified carrot reference material (NCSZC73031, China National Analysis Center for Iron & Steel, Beijing, China) was integrated in every set of 24 beakers. Four ml of nitric acid (65 %) was added to every sample. The samples were heated in an oven (VO400, manufactured by Memmert, Schwabach, Germany) for 7 h at 180°C. The cooled samples were filled up with deionized water to a volume of 25 ml and filtered through ashless filter paper (MN640 m, Ø125 mm, manufactured by Macherey-Nagel, Düren, Germany). Absorption was measured at 213.8 nm for Zn and 248.3 nm for Fe with an atomic absorption spectrometer (model 11003, manufactured by PerkinElmer, Waltham, MA, USA). A blank was subtracted from the measured values and the mineral concentration of every sample was calculated by using fitting standard solutions.

For the phytate/mineral molar ratios, the measured concentrations were divided by their molar mass (phytate 660.04 g mole^−1^; Fe 55,845 g mole^−1^; Zn 65.38 g mole^−1^).

### Data analysis

Statistical analyses were conducted using the statistical software SAS 9.4 (SAS institute Inc., Cary, NC, USA). For all parameters the general linear model (GLM procedure) with least square means and Tukey-Kramer adjustment for multiple comparisons was calculated for the factors genotype, treatments and their interaction. For selected traits a comparison between the two groups ‘cultivated genotypes’ (*O. sativa* and *O. glaberrima*) and ‘wild genotypes’ (all non-cultivated species) in the different treatments was undertaken by two-way analysis of variance (ANOVA).

## Results

Plants treated with Fe started to develop foliar symptoms of Fe toxicity soon after the first Fe application six and seven WAT. In the acute stress treatment, the peak of symptom formation was reached at eight WAT, while in the chronic stress treatment, the average symptom level continued to rise until 14 WAT (Table 1). From eleven WAT, the average symptom level was significantly higher in the chronic stress treatment compared to the acute stress treatment (Table 1). Significant genotypic differences in symptom formation occurred on all sampling days (Table 1) and were reflected in a broad range of responses of individual genotypes in terms of average LBS across all sampling days (Figure 1). In both stress conditions, the most sensitive genotypes were RWR, but on the other hand, some RWR also ranked among the most tolerant genotypes, especially in the acute Fe stress. Here, the two most tolerant accessions were from the species *O. meridionalis* (Figure 1a). On the other hand, in the chronic Fe stress, the most tolerant genotypes were *O. sativas*. These had previously been described as Fe tolerant *i.e.* FL483 [26] and Kitrana 508 [32]. Apart from that, there was no clear pattern regarding the tolerance of domesticated and wild rice species, as both groups exhibited a broad range of differential responses.

**Table 1:**
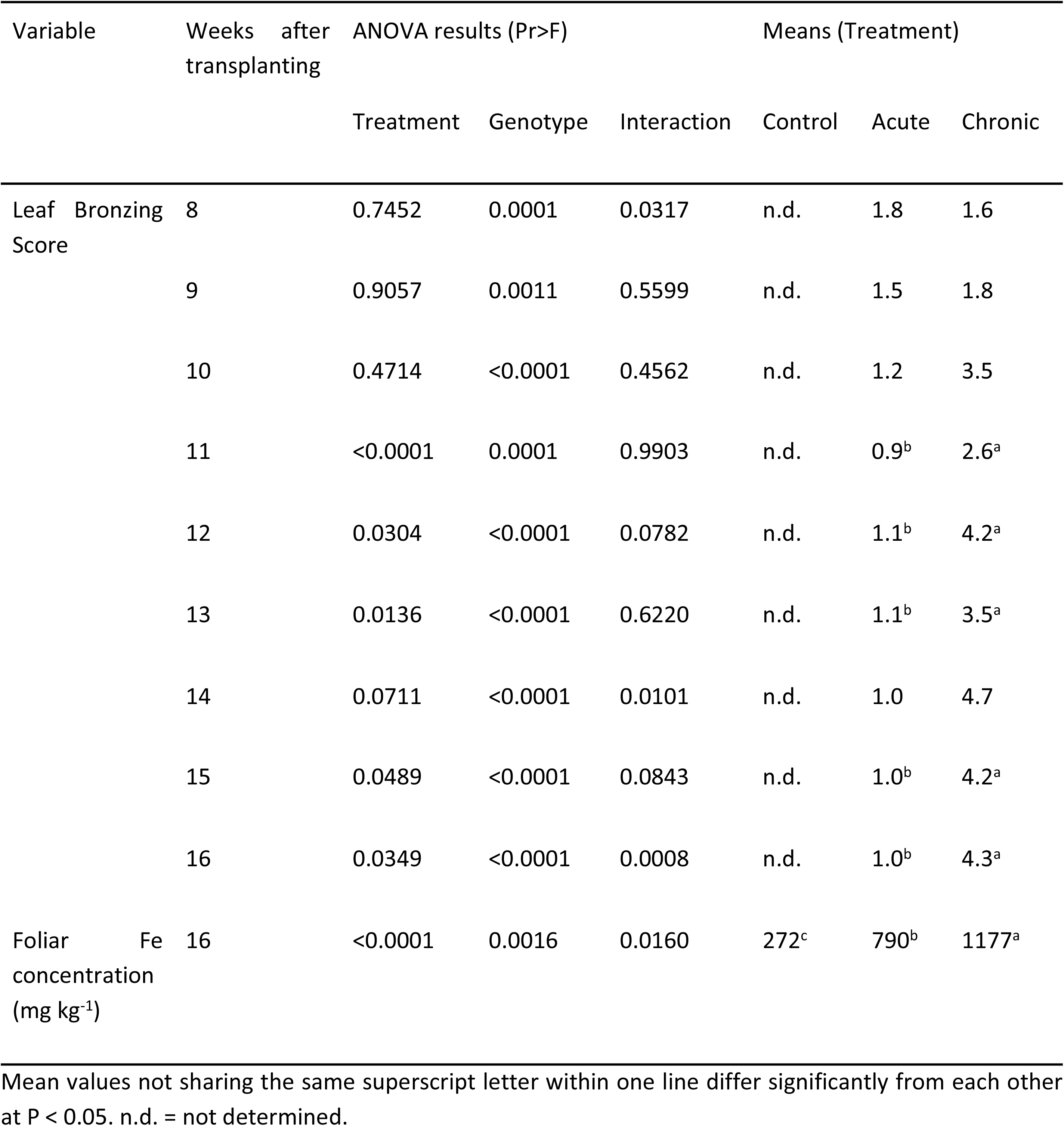
ANOVA results and treatment mean values for traits determined during the vegetative growth of wild and domesticated rice species exposed to acute or chronic iron toxicity

**Figure 1:**
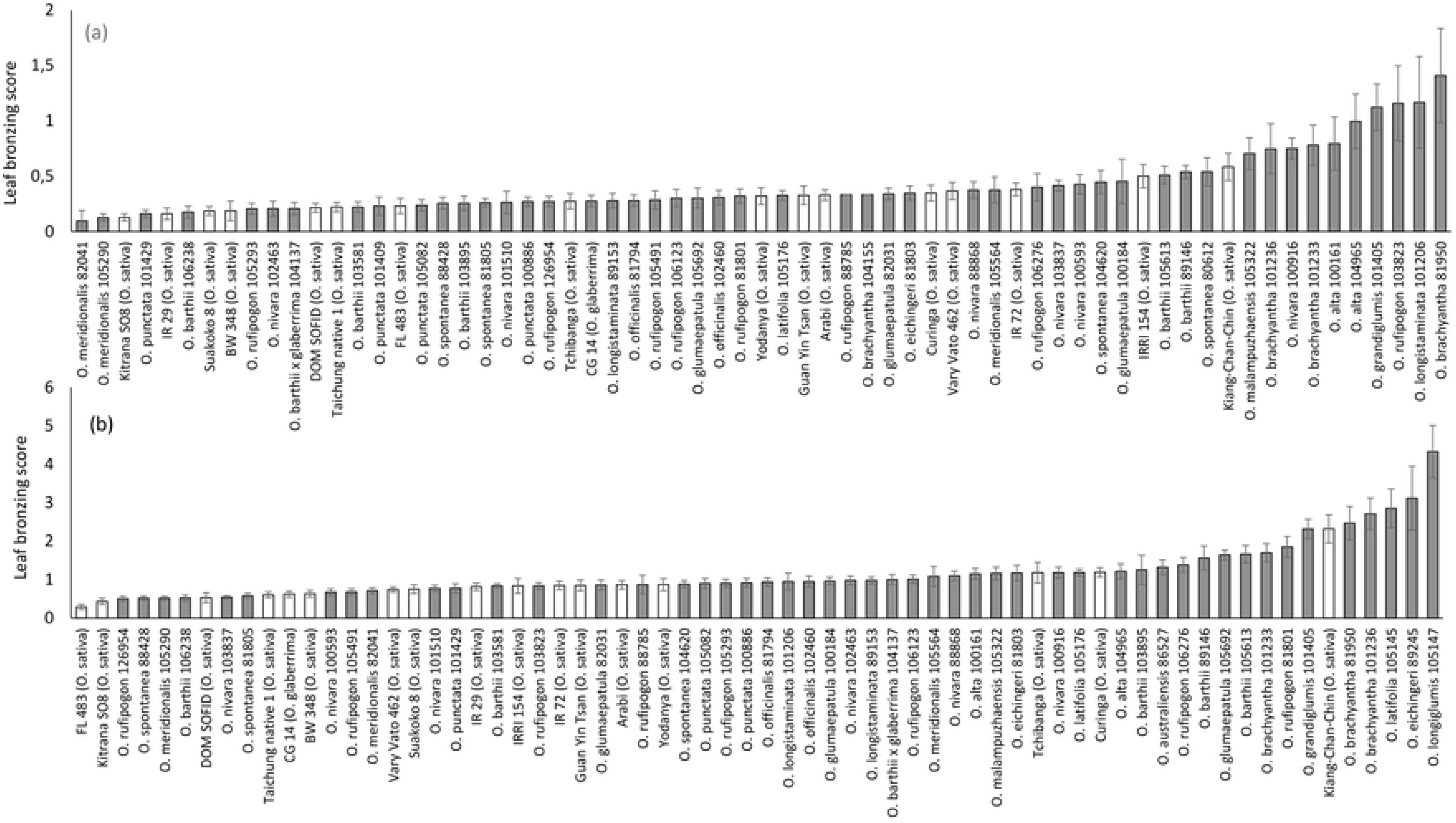
Leaf bronzing score of domesticated rice varieties and rice wild relatives in acute (a) and chronic (b) Fe toxicity stress. Mean values and standard errors across nine sampling days, two experimental and four sub-replicates are plotted. White bars represent domesticated varieties while grey bars represent wild relatives

Foliar Fe analyses of selected genotype at 16 WAT demonstrated that even in the acute Fe stress, where Fe was applied already six and seven WAT, most genotypes had values exceeding 300 mg kg^−1^, which is widely accepted as the threshold for Fe toxicity [35]. Average values were significantly higher in the chronic stress treatment compared to the acute stress (Table 1). While the highest Fe levels in control conditions occurred in *O. sativas* (Fig. 2a), the opposite was seen in chronic Fe stress (Fig. 2c). In contrast, RWR accessions had the lowest Fe concentrations in all three conditions, suggesting efficient Fe exclusion mechanisms.

**Figure 2:**
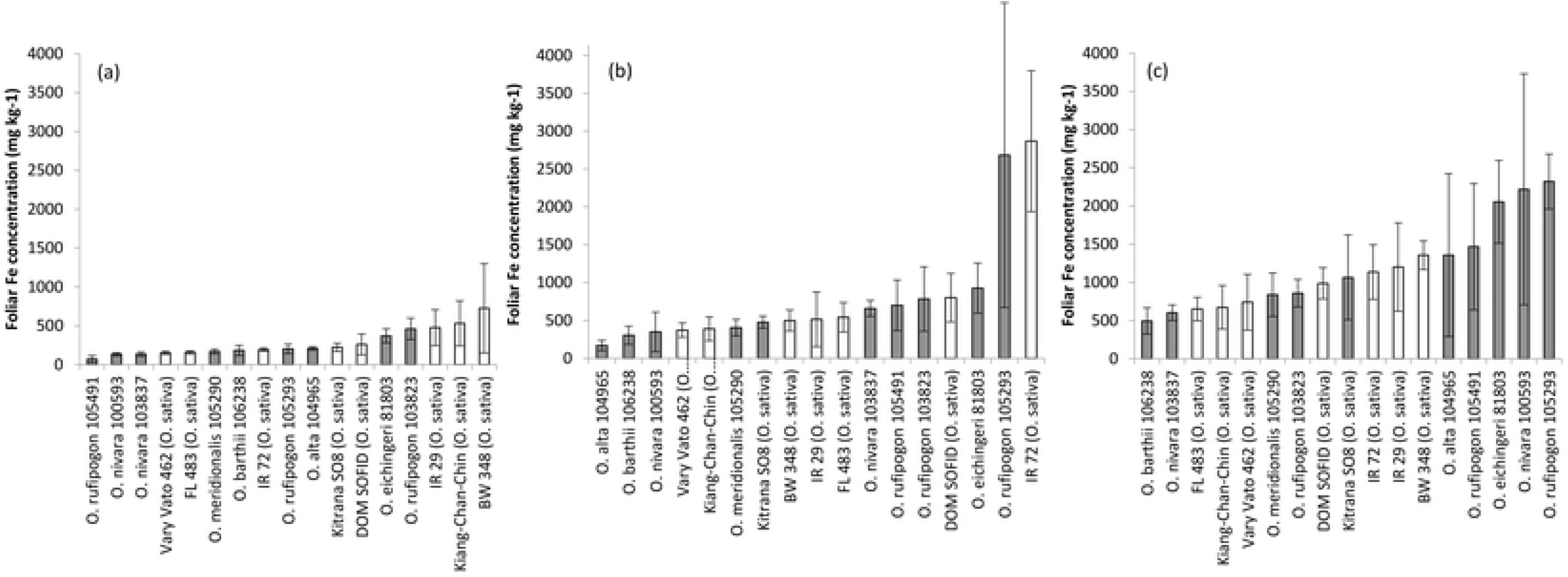
Foliar Fe concentrations of selected domesticated rice varieties and rice wild relatives in control conditions (a) acute (b) and chronic (c) Fe toxicity stress. Mean values and standard errors are plotted (n= 3-9). White bars represent domesticated varieties while grey bars represent wild relatives.

At harvest we determined shoot morphology and straw biomass of all plants, and analyzed grain yield and quality data for those accessions, in which a sufficient number of replicated plants produced seeds. This was true for most of the domesticated rice accessions, but only ten RWR. The Fe stress treatments did not negatively affect straw biomass and plant morphological traits, in which a rather positive effect was seen especially in the acute Fe treatment (Table 1). In contrast, grain yield was negatively affected in the chronic Fe treatment, but not in the acute Fe treatment (Table 2). In both Fe treatments, the sterility rate and the harvest index were significantly negatively affected compared to the control (Table 2). Analyses of the grain yield responses of individual genotypes revealed that most accessions had slightly enhanced grain yield in the acute Fe treatment (Figure 3a). In contrast, grain yield was negatively affected in most accessions in the chronic Fe treatment, except for two *O. sativas* (Dom Sofid and Kitrana 508) and one RWR (*O. rufipogon*), which had the highest grain yield among all accession that produced seeds. As opposed to the cultivated rice accessions, the group of RWR did not respond to Fe treatment with an increase in sterility (Figure 4).

**Table 2:** ANOVA results and treatment mean values for yield and grain quality traits of wild and domesticated rice species exposed to acute or chronic iron toxicity

| Variable | ANOVA results ($Pr > F$) | | | Means (Treatment) | | |
| --- | --- | --- | --- | --- | --- | --- |
|  | Treatment | Genotype | Interaction | Control | Acute | Chronic |
| *Straw yield (g plant<sup>-1</sup>)* | <0.0001 | <0.0001 | 0.029 | 54.7<sup>b</sup> | 70.5<sup>a</sup> | 59.2<sup>b</sup> |
| *Plant height (cm)* | <0.0001 | <0.0001 | <0.0001 | 167.6<sup>b</sup> | 177.4<sup>a</sup> | 176.6<sup>a</sup> |
| *Tiller number* | <0.0001 | <0.0001 | 0.0116 | 12.0<sup>b</sup> | 14.95<sup>a</sup> | 12.6<sup>b</sup> |
| *Grain yield (g plant<sup>-1</sup>)* | <0.0001 | <0.0001 | 0.8394 | 18.9<sup>b</sup> | 23.6<sup>a</sup> | 15.2<sup>c</sup> |
| *Panicle number* | <0.0001 | <0.0001 | 0.2228 | 9.7<sup>b</sup> | 12.8<sup>a</sup> | 10.6<sup>b</sup> |
| *Seeds per panicle* | 0.9189 | <0.0001 | <0.0001 | 132.5 | 131.3 | 129.2 |
| *Spikelet sterility (%)* | <0.0001 | <0.0001 | <0.0001 | 50.1<sup>a</sup> | 55.9<sup>b</sup> | 62.2<sup>c</sup> |
| *Thousand kernel weight (g)* | <0.0001 | <0.0001 | <0.0001 | 20.8<sup>a</sup> | 20.4<sup>a</sup> | 19.3<sup>b</sup> |
| *Harvest Index (%)* | <0.0001 | <0.0001 | 0.5422 | 27.9<sup>a</sup> | 23.3<sup>b</sup> | 18.4<sup>c</sup> |
| *Grain Fe concentration (mg kg<sup>-1</sup>)* | 0.2584 | 0.0299 | 0.1963 | 31.5 | 32.3 | 39.0 |
| *Grain Zn concentration (mg kg<sup>-1</sup>)* | <0.0001 | <0.0001 | <0.0001 | 25.6<sup>b</sup> | 28.3<sup>ab</sup> | 30.8<sup>a</sup> |
| *Phytate (mg g<sup>-1</sup>)* | 0.007 | <0.0001 | <0.0001 | 9.7<sup>a</sup> | 10.4<sup>b</sup> | 9.6<sup>a</sup> |
| *Phytate/Fe molar ratio* | 0.14 | 0.0197 | 0.234 | 40.4 | 39.5 | 29.7 |
| *Phytate/Zn molar ratio* | 0.007 | <0.0001 | 0.0003 | 38.0<sup>b</sup> | 36.3<sup>b</sup> | 31.6<sup>a</sup> |

Treatment means with different superscript letters in the same line indicate significant difference ( $p < 0.05$ ).

**Figure 3:**
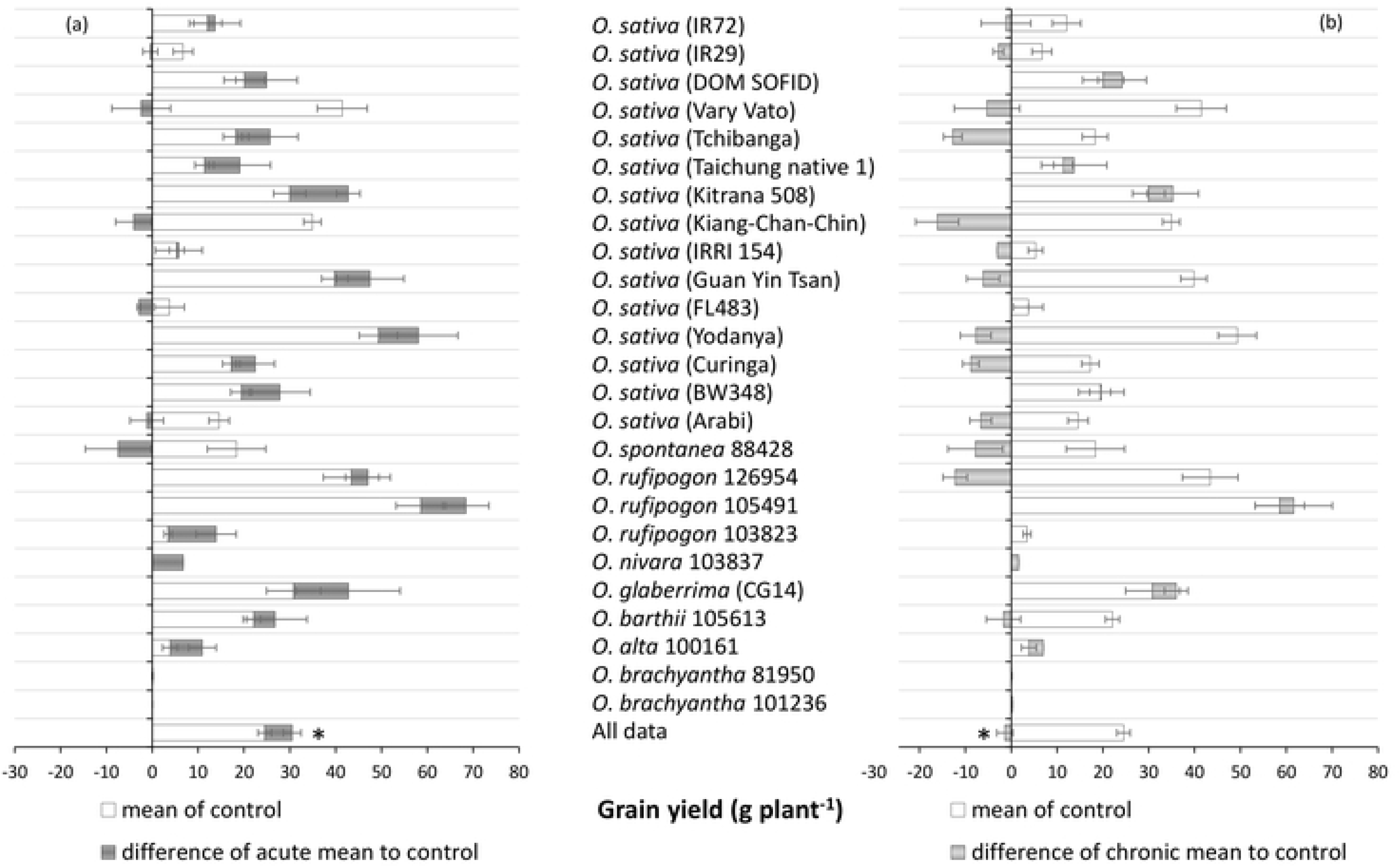
Grain yields of selected domesticated rice varieties and rice wild relatives in acute (a) and chronic (b) Fe toxicity stress compared to control conditions. The grey bars indicate the negative (directed towards the left) or positive (directed towards the right) difference from the control. Mean values and standard errors (n=2-8) are plotted. Asterisk indicates a significant difference from the control at P<0.05.

**Figure 4:**
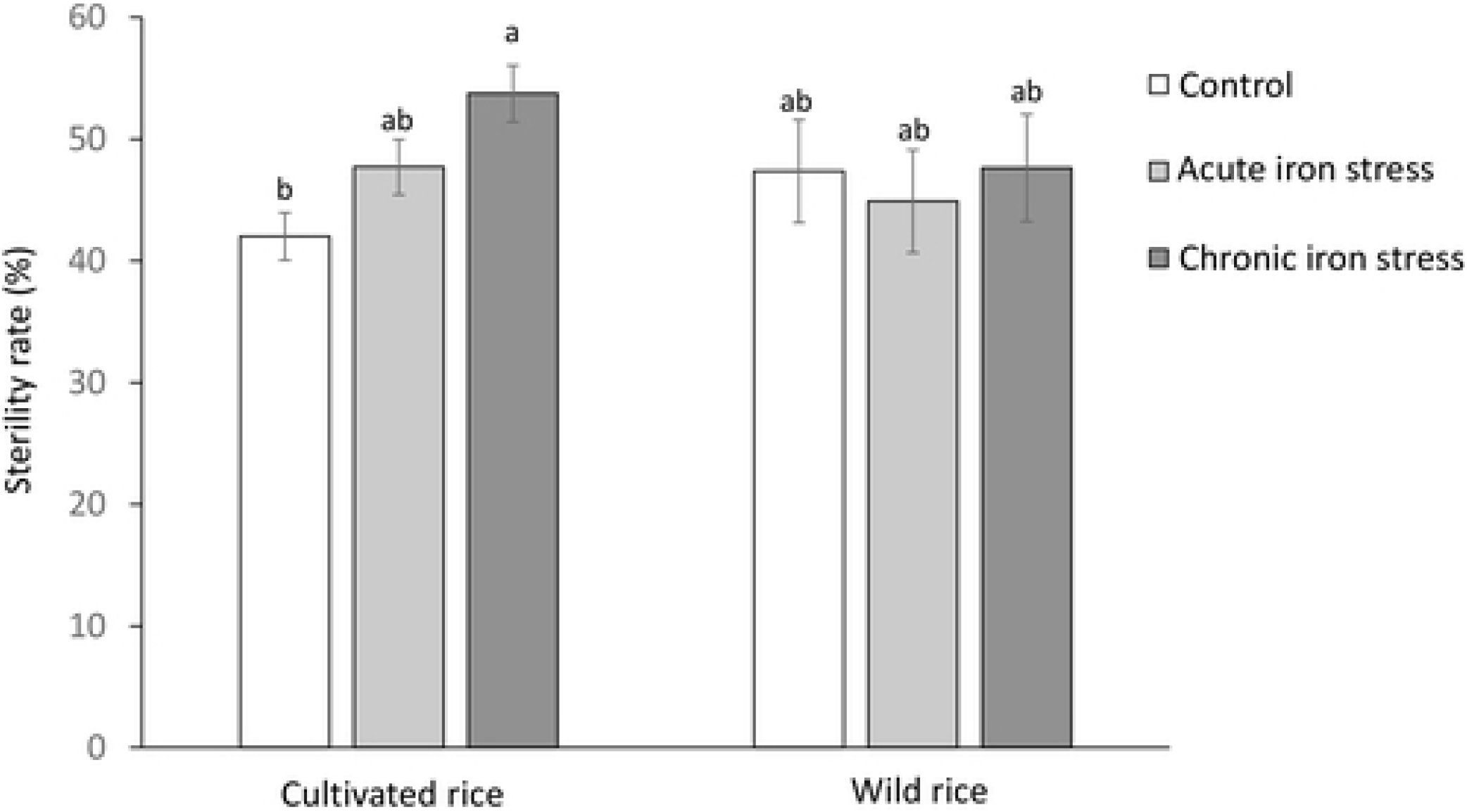
Sterility rates of cultivated and wild rice species in control conditions, acute and chronic Fe toxicity. Mean values and standard errors are plotted (n=33-115). Bars not sharing the same letter differ from each other at P<0.05 by Tukey’s HSD test.

Grain Fe concentrations were slightly elevated in the Fe treatment but the differences from the control were not significant (Table 2). In contrast, grains produced in the chronic Fe treatment had elevated Zn concentration and lower phytate /Zn molar ratio, and grains produced in the acute Fe treatment had elevated phytate concentrations (Table 2). Comparison of grain quality traits in selected accessions revealed that the *O. sativa* Dom Sofid had the highest Fe concentration, whereas two RWR showed the highest Zn levels (Fig. 5b), which were also associated with high phytate levels (Figure 5c). Very low phytate level occurred in an *O. spontanea* accession.

**Figure 5:**
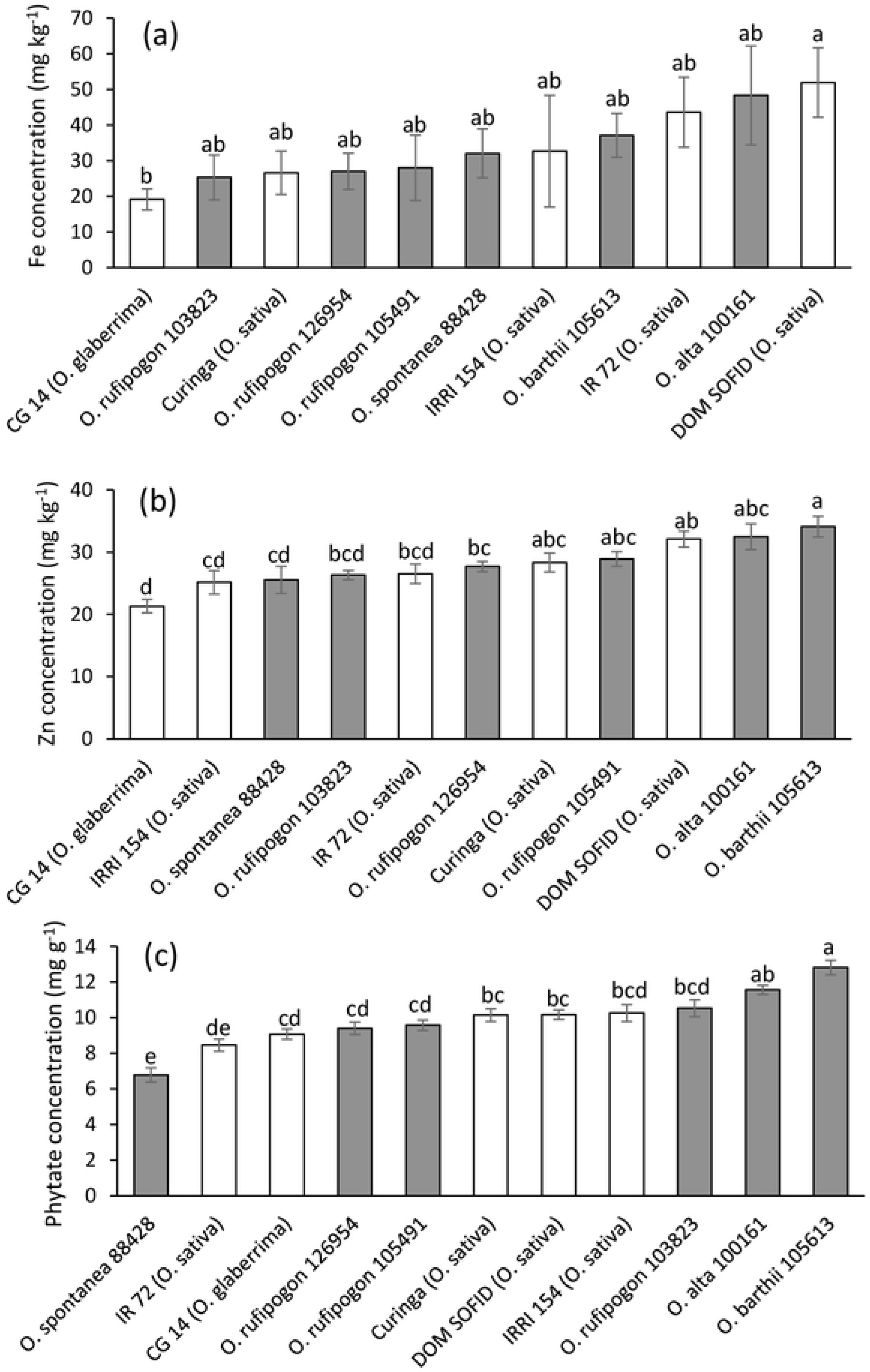
Brown rice mineral concentrations of selected domesticated rice varieties and rice wild relatives averaged across different treatments. (a) Fe concentration; (b) Zn concentration; (c) Phytate concentration. Mean values and standard errors (n=5-12) are plotted. Bar not sharing the same letters differ significantly from each other by Tukey HSD test.

## Discussion

The treatments in the present study (acute and chronic Fe stress) were conceived in order to simulate two types of Fe stress typically occurring in the field. Acute Fe toxicity frequently occurs in inland valleys during heavy rainfall events, leading to a transient peak of interflow of reduced Fe from adjacent slopes into the rice field. This type of Fe stress is wide-spread for example in West Africa [23]. On the other hand, chronic Fe stress occurs on naturally Fe rich soils in which Fe stress builds up more gradually during the growing season upon flooding. This type of stress often occurs on acid sulfate soils or Ferralsols and is widespread in Southeast Asia, the South of Brazil or Madagascar [23]. The semi-artificial experimental setup used in this study was tested previously with six *Oryza sativa* varieties differing in Fe tolerance [28]. Similar to that previous study, acute Fe stress lead to the development of stress symptoms, but did not negatively affect biomass and grain yields (Table 2). In contrast, stress symptoms were more pronounced in chronic Fe stress, which also led to significant reductions in grain yield (Table 2), largely due to higher spikelet sterility (Figure 4). Therefore, our study confirms previous observations that plants respond differently to chronic and acute Fe stress on a physiological level [36], or in terms of yield [28], and that plants can compensate for early Fe tress comparatively well. The phenomenon of positive growth responses to mild metal stresses has been termed as ‘hormesis’ [37], and may occur due to physiological processes including the activation of antioxidants and complex signaling pathways. In contrast, the grain yield losses due to enhanced sterility in the chronic Fe treatment may be a consequence of oxidative stress in reproductive organs. High levels of Fe lead to the Fenton reaction, in which ferrous Fe reacts with hydrogen peroxide to form ferric Fe, hydroxide and the hydroxyl radical [25,38]. The hydroxyl radical is an extremely harmful molecule which cannot be effectively scavenged by the plants’ antioxidant system [39].

Genetic variation in adaptive traits is the prerequisite for the breeding of new varieties with enhanced tolerance to Fe toxicity. In this study, both domesticated and wild rice species showed substantial variation in all observed traits. Foliar stress symptoms are often used as a secondary trait indicating Fe tolerance, as it has shown relatively good correlation with important agronomical traits such as grain or biomass yield in field experiments [27], and they can be scored for a large number of plants in comprehensive screening experiment. For this trait, the most sensitive genotypes in both stress conditions were all RWR (Figure 1). This high level of Fe sensitivity compared to domesticated rice may be explained with the fact that RWR often grow in non-permanently flooded soils, therefore lacking adaptive mechanisms such as Fe exclusion, which becomes relevant in permanently flooded conditions, where soluble Fe is abundantly available. Rice domestication and breeding has been accompanied by the development of irrigation infrastructure and the permanent flooding of rice soils, making adaptation to high amounts of soluble Fe a trait that breeders may unintentionally have selected for in cultivated rice varieties. Nevertheless, extremely low LBS of some RWR, especially *O. meridionalis* accessions (Figure 1a) indicate that specific RWR do possess adaptive genes which can supplement the gene pool for adaptive rice breeding. The high level of adaptation of this particular species may have an evolutionary background as it is often found at the edges of freshwater lagoons, temporary pools and swamps of the Northern territories of Australia [40], where Fe-rich soils such as Ferrosols occur [41]. As a member of the *O. sativa* complex with an AA genome, *O. meridionalis* represents an easily accessible gene pool, and crosses with *O. sativa* have previously been reported [31].

Foliar Fe levels in both Fe stress treatments were mostly above 300 mg kg^−1^, which is considered to be the threshold for Fe toxicity [35]. However, one *O. alta* accession maintained very low foliar Fe level in the acute Fe stress (Figure 2b). It is doubtful whether this occurred due to an active Fe exclusion mechanism such as Fe oxidation in the rhizosphere [26,42]. More likely, Fe was ‘diluted’ as this species reached an average height of 316 cm and thus produced a lot of biomass for each of its few tillers (3.7 tillers on average). This plant architecture is not desirable in rice breeding, thus *O. alta* may not be a suitable donor of Fe tolerance traits. The low Fe levels observed in two RWR (*O. barthii* and *O. nivara*) in chronic Fe stress (Figure 2c) appear more promising, especially as these two species form part of the AA genome group with lower crossing barriers towards *O. sativa* or *O. glaberrima*.

The analysis of grain yields was to some extent compromised by the fact that many RWR did not set seeds within the experimental period. Factors contributing to this phenomenon include long vegetation cycles, specific light and day length requirements, and self-incompatibility of many of the RWR [40]. Although experimental conditions including climate were controlled within a range that supports the growth of most *Oryza* species, the large heterogeneity of screened genotypes is difficult to compensate or adequately address by standardized climatic conditions. Among the ten RWR that produced seeds, four showed higher yields in the chronic Fe stress than in the control, even though the overall yield response across all screened genotypes was significantly negative in this treatment (Figure 3). This positive yield response was not unique to RWR, as it also occurred in some *O. sativa* accession that had previously been described as Fe tolerant [28,32], but the overall highest grain yield was seen in an *O. rufipogon* accession (Figure 3b), and increases in spikelet sterility due to chronic Fe stress occurred in cultivated rice varieties rather than RWR (Figure 4).

The concentrations of essential mineral nutrients in rice grains are a matter of concern because nutritional deficiencies in Fe and Zn are very widespread [43,44]. Enriching rice grains in these elements (‘biofortification’ [45]) is therefore seen as one out of several possible strategies to combat mineral disorders in human diets. On the other hand, low phytate levels in cereal grains are desirable from a nutritional perspective as this P-storage compound forms insoluble complexes with Fe and Zn thereby limiting the bioavailability of these elements in human diets [46]. Our present study confirms previous findings in domesticated rice varieties [28] that growing rice in high Fe conditions does not significantly increase grain Fe concentrations in rice grains of most genotypes (Table 2), despite substantial increases in Fe concentrations in vegetative tissue (Figure 2). This indicates that rice plants possess physiological barriers protecting reproductive tissue from Fe levels that could cause oxidative stress and consequently sterility. The regulation of metal transporters responsible for the loading of metals from mother plant tissues such as seed coats into filial tissues [47] could be such a mechanism. On the other hand, average Zn concentration was enhanced in plants grown in chronic Fe stress (Table 2), which could be due to a concentration effect, which occurs when plants grown under stress form less biomass or yield with the same amount of available mineral elements[48]. Irrespective of the treatment and genotype, phytate/Fe and phytate/Zn molar ratios were in a range indicating relatively poor bioavailability [49], which is characteristic of unrefined cereals evaluated individually, and not as part of a diverse diet. While the RWR did not stand out with extraordinarily high nutritional value, positive features were seen especially in terms of high Zn concentration (Figure 5b) and low phytate concentration (Figure 5c).

## Conclusion

We screened a broad range of RWR for adaptive and grain quality traits when grown in Fe toxic conditions, and compared them to domesticated varieties that had been pre-selected to include some with a previously-known high level of tolerance. None of the RWR drastically outperformed the domesticated varieties, which is partly due to their poor agronomic traits. Nevertheless, RWR exhibited promising performance in individual traits such as low symptom formation (*O. meridionalis*), high grain yield (*O. rufipogon*) or low grain phytate (*O. spontanea*). Chromosomal segment substitution lines of interspecific crosses in *O. sativa* background [31] are available, facilitating future in-depth physiological analyses in more uniform screening experiments due to lower morphological and phenological diversity of such crosses as compared to the very diverse material screened in this study. The availability of genome sequences of RWR [5] provides an additional powerful resources for the discovery of genes underlying some of the adaptive traits identified in this study. Therefore, this study provided an important first step in expanding the gene pool towards RWR for the breeding of rice varieties adapted to Fe toxicity.

## Acknowledgements

This study was financially supported by the Foundation *fiat panis*. The authors wish to thank the gene bank of the International Rice Research Institute (IRRI) for providing seeds, and Dr. Hannes Dempewolf from the Global Crop Diversity Trust in Bonn (Germany) for inspiring this study.

## Supporting Information

Supplementary Table S1: List of germplasm used in the screening experiment

